# Codon bias, nucleotide selection, and genome size predict *in situ* bacterial growth rate and transcription in rewetted soil

**DOI:** 10.1101/2024.06.28.601247

**Authors:** Peter F. Chuckran, Katerina Estera-Molina, Alexa M. Nicolas, Ella T. Sieradzki, Paul Dijkstra, Mary K. Firestone, Jennifer Pett-Ridge, Steven J. Blazewicz

**Affiliations:** Department of Environmental Science, Policy, and Management, University of California, Berkeley, CA, USA; Physical and Life Sciences Directorate, Lawrence Livermore National Laboratory, Livermore, CA, USA; Department of Plant and Microbial Biology, University of California, Berkeley, CA, USA; Department of Biological Sciences, Columbia University, NY, NY, USA; Laboratoire Ampère, École Centrale de Lyon, Lyon, France; Center for Ecosystem Science and Society (ECOSS) and Department of Biological Sciences, Northern Arizona University, Flagstaff, AZ, USA; Life & Environmental Sciences Department, University of California Merced, Merced, CA, USA; Innovative Genomics Institute, University of California Berkeley, Berkeley, CA, USA

## Abstract

In soils, the first rain after a prolonged dry period greatly impacts soil microbial community function, yet we lack a full understanding of the genomic traits associated with the microbial response to rewetting. Genomic traits such as codon usage bias and genome size have been linked to bacterial growth in soils—however this is often through measurements in culture. Here, we used metagenome-assembled genomes in combination with metatranscriptomics and ^18^O- water stable isotope probing to track genomic traits associated with transcriptional activity and growth of soil microorganisms over the course of one week following rewetting of a grassland soil. We found that the codon bias in ribosomal protein genes was the strongest predictor of growth rate. We also observed higher growth rates in bacteria with smaller genomes, demonstrating that reduced genome size contributes to bacterial growth responses to sudden changes in water or nutrient availability—potentially explaining why smaller genomes are more prevalent in arid and carbon poor systems. High levels of codon bias corresponded to faster transcriptional upregulation of ribosomal protein genes. In early transcribing taxa, nucleotides requiring less energy to produce were more common at synonymous substitution sites—where nucleotide substitutions did not change the encoded amino acid. We found several of these relationships also existed within a phylum, suggesting that association between genomic traits and activity could be a generalized characteristic of soil bacteria. These results provide *in situ* evidence that following rewetting, certain genomic characteristics affect soil microbial growth rate and transcription, and points towards the fitness advantages that these traits might pose for bacteria under changing conditions in soil.

## INTRODUCTION

Rewetting is an ecologically important soil phenomenon where microbial traits associated with growth and activity may play an important role in biogeochemical cycling, and community structure and function. In ecosystems characterized by dry and wet seasons, the first rain event following the dry season results in a large pulse of CO_2_ (known as the Birch Effect) (1), which can account for a large portion of annual soil C loss (2). During these events, rapid stimulation of microbial activity and growth is driven by both the release of water stress as well as an influx of bioavailable carbon compounds sourced from osmolytes (3–5), microbial necromass (6, 7), slaking of microaggregates (8), and increased connectivity in the soil matrix (9, 10). These microbial responses to rewetting therefore have important consequences for nutrient cycling and ecosystem function. However, the microbial traits that underly the growth and activity during these events remain unknown.

Trait-based frameworks have long-been used in ecology to help understand complex community patterns (11, 12), and this approach has gained considerable attention in microbial ecology (13, 14), where advances in sequencing technology have expanded the ability to probe microbial attributes. Analysis of bacterial traits enables two key insights into soils. First, since traits are the product of evolutionary forces, understanding the distribution of traits progresses fundamental questions concerning the evolution and selection for certain attributes (15). Second, considering the complexity of microbial community composition, traits serve as valuable metrics for assessing community dynamics across scales (16, 17) and with ecosystem function (14).

Traits associated with growth rate are of particular interest, as growth rate is often central in life-strategy frameworks (14, 18, 19), functional metrics such as carbon use efficiency (20, 21), and understanding changing community dynamics.

Trait-based analyses using sequence data can generally be separated into two broad categories: functional traits (those which encode for specific cellular functions) and genomic traits (characteristics of a genome that do not describe taxonomy, anatomy or a specific function, e.g., GC-%, genome size, codon usage, etc.). Functional traits are often evaluated based on functional gene sequences and their expression in different environmental conditions. During growth, expression of specific genes varies considerably depending on function (22) and environmental constraints (23). Several functional genes have been associated with growth in soil microbial communities (24), but the gene-growth relationship is often tied to a specific set of conditions (e.g., favoring specific nutrient acquisition strategies or metabolic pathways) and may not be predictive outside of that environment. Outside of ribosomal protein genes, which are reliably upregulated during growth, there are relatively few genes that have been universally tied to growth. The number of rRNA gene copies in a genome positively correlates with maximum growth rate (25, 26)—making rRNA gene copies a useful trait dimension used in microbial ecology trait-based frameworks (18, 19, 27). However, even the link between rRNA levels and growth/activity is inconsistent (28), highlighting the limitations of using functional genes and gene transcription as predictive traits of growth.

In contrast, genomic traits (such as genome size, nucleotide frequency, and codon usage) may provide comprehensive metrics that are predictive of activity and growth (26, 29). For example, transcription and growth rate may be strongly influenced by the frequency with which certain codons occur in a gene sequence. Generally, this occurs when codon frequency aligns with the tRNA pool (30). Multiple codons can encode for the same amino acid, and these “synonymous codons” have a greater affinity for tRNA with corresponding anticodons. When the synonymous codons of a gene sequence occur in a similar frequency to the pool of tRNA anticodons, this can increase the rate of both transcription and translation—and the degree of this alignment is referred to as codon optimization (30). For translation, codon optimization impacts the rate of elongation, protein folding, initiation, and termination (31–33). For transcription, codon optimization often predicts mRNA abundance since more optimized codons generally increases mRNA stability (34, 35) and is related to higher levels of transcription (36, 37). Codon bias—the degree of redundancy of codon usage in the genetic code for a particular gene or genome—correlates with the degree of codon optimization (38). High levels of codon bias in ribosomal proteins is associated with rapid growth in bacteria (26, 39) and has increasingly been used to predict growth rate (29, 40).

Outside of an intrinsic relationship between GC content and codon usage (where very high or low GC content tend to naturally have high levels of codon bias) (30), there appears to be no clear relationship between doubling time and GC content in bacteria (41). However, the cost of nucleotide synthesis does influence transcription, and rapidly transcribed genes often use less energy expensive (‘cheaper’) nucleotides at synonymous sites to reduce the cost of transcription (42). Using metabolic models of *E. coli*, Chen et al. 2014 (42) estimated that guanine requires 4.6 more ATP to produce than cytosine, and adenine requires 7.7 more ATP to produce than uracil. Therefore, genes with cytosine and thymine at synonymous sites tend to be transcribed faster than genes with guanine and adenine (42). At nonsynonymous sites (where a nucleotide substitution changes the encoded amino acid), there is an inverse relationship between nucleotide cost and the cost of their encoded amino acid sites (42)—such that a higher frequency of more expensive nucleotides at nonsynonymous sites is associated with higher levels of expression.

Genome size is also thought to influence growth rate; however, the relationship is less clear than codon usage or rRNA copy number. In oligotrophic marine environments, extremely small genomes are thought to arise in response to nutrient limitation to curb the cost of reproduction (termed genomic streamlining; (43)), but may have slower maximum growth rates than copiotrophs with larger genomes (18). Although streamlined genomes have been documented in soils (44–46), soil bacterial genomes tend to be large relative to other ecosystems (47)— potentially because of the increased metabolic diversity required to utilize complex substrates (48). It has been hypothesized that these large genomes might come at the expense of growth rate (49), as the increased energy required for reproduction in large genomes may slow growth. However, the evidence regarding the relationship between genome size and growth rate in soil bacteria is inconclusive (50, 51), necessitating further investigation.

In soils, a major hurdle in assessing how genomic traits relate to growth rate has been our inability to effectively measure these traits *in situ*. Much of the work linking these traits to growth has been conducted using soil isolates where growth rates are assessed in pure culture experiments; however, traits associated with media-based growth are not reliable predictors for microbial growth in a natural environment (51). Recent method and technical advances have created new opportunities to track microbial growth *in situ*. In quantitative stable isotope probing (qSIP), the incorporation of added stable isotopes into DNA tracks the growth rate of microbes (52). This approach has enabled valuable insights into metagenomic features and metagenome-assembled genomes (MAGs) associated with growth (7, 24, 53, 54), and shows promise for linking measurable microbial traits and growth rates *in situ*. In this study, we combined metagenomic, metatranscriptomic, and qSIP data to assess how genomic traits correspond with activity and growth during the rewetting of seasonally dry soil in a Mediterranean grassland. Specifically, we investigated how traits such as genome size, nucleotide selection, codon usage, and ribosomal protein nucleotide and codon frequency relate to growth and transcription. We hypothesized that the bacterial response to wet-up would be associated the following set of genomic traits: 1) codon bias in ribosomal protein genes would be a strong predictor of both transcription and growth rate, as based on the well-established relationship between codon bias and growth (29); 2) fast-growing and transcribing bacteria would use biosynthetically cheaper nucleotides (i.e. C and T/U, as opposed to G and A) at synonymous substitution sites, and; 3) genome size would have little impact on growth, as previous studies have shown factors such as genome and cell size do not strongly relate to growth in soil bacteria (50, 51). Through a combined multi-omics and qSIP approach, we aim to test these hypotheses and gain a better understanding of the mechanisms which affect this fundamental response in soils.

## METHODS

The field study was located at Hopland Extension and Research Center (HERC), Hopland, California, USA (39° 00 14.6 N, 123° 05 09.1 W), which resides on the ancestral home of the Shóqowa and Hopland people. The region features a Mediterranean climate of warm, dry summers and cool, wet winters. *Avena barbata* (wild oat grass) dominated the studied field site. This study consisted of 16 plots 3.24 m^2^, with rainout shelters constructed around each plot, either allowing full or 50% mean annual precipitation for the two preceding years. The soils are of the Squawrock-Witherell complex, with pH of 7.3, total C content of 15.1 mg/g and total N of 1.5 mg/g. A full description of the field experimental set-up can be found in Fossum et al. 2022 (55).

Soils were collected in September 2018 at the end of the dry season and 25 days before the first rainfall of the wet season. At the time of collection, the soil gravimetric water content was approximately 3%. Topsoil samples (0-15 cm deep, roughly 4700 cm^3^) were taken from eight plots (four full and four reduced precipitation). Samples were transported to Lawrence Livermore National Laboratory where soil from each field plot was separately homogenized and sieved (2 mm) to remove large rocks and roots.

### Wet-up experiment

Details of the soil H ^18^O labeling can be found in Nicolas et al. 2023 and Sieradzki et al. 2022 (7, 56). In brief, soils from each of the eight plots were separated into 11 microcosms containing 5 g of soil each, for 88 samples in total. Soils were brought to 22% gravimetric water content by adding 1 ml of either natural abundance water or 98 atom % H_2_^18^O, and then incubated in the dark within sealed 500 ml mason jars. Four replicates of each sample type (^18^O labeled and unlabeled) from each plot were destructively harvested at 3, 24, 48, 72, and 168 h post wet-up— in addition to a set of dry soil controls (i.e. 0 h). Samples were immediately frozen in liquid-N_2_ and stored in a freezer at −80 °C.

### Metagenomic qSIP

A full description of the metagenome DNA extraction, sequencing, assembly, binning, and qSIP calculations is provided in Sieradzki et al., 2022. Briefly, DNA from three plots per treatment and timepoint was extracted in triplicate using a phenol chloroform extraction protocol adapted from Barnard et al. (2015) (57) and then pooled. 5 µg DNA samples were then spun at 20 °C for 108 hours (176,284 RCF_avg_) on a Beckman Coulter Optima XE-90 ultracentrifuge in a cesium chloride solution (density of 1.730 g mL^-1^) to create a density gradient following a technique previously described in Blazewicz et al. 2020 (58). The contents of the ultracentrifuge tube were then separated into 36 fractions using Lawrence Livermore National Laboratory’s high-throughput SIP pipeline (54), each of which was assessed for density and DNA concentration. Fractions were combined into five groups along the density gradient, purified and concentrated, and then sequenced on an Illumina Novaseq S4 2x150bp platform. Adapters were trimmed and reads were QC filtered using bbduk (59) and Sickle (60). QC filtered reads were assembled into contigs using MEGAHIT (v1.2.9; (61)), binned with MaxBin 2.0 (62) and MetaBAT2 (63), and refined with MetaWrap (64).

We combined this set of MAGs with another set generated from a study at a nearby site at HREC (24) and dereplicated the combined set of MAGs using dRep, which also assesses MAG completeness (65). Open reading frames were identified using Prodigal (66) and annotated with KEGG Orthologs (67) using DRAM (68).

Reads from each SIP fraction were then mapped against the combined set of MAGs using BBMap (59). The mapping of reads from each fraction was used to calculate atom fraction excess (AFE) for each MAG (described in ref. 56), which quantitates the amount of isotopic label incorporated into a genome. Since reproducing genomes incorporate the ^18^O isotope from the added ^18^O-water into their DNA, AFE values can be used as an index of growth (52).

### RNA Extraction and sequencing

RNA was extracted from four soil samples from each time-point and precipitation treatment using the RNeasy PowerSoil Total RNA kit (Qiagen) according to manufacturer instructions. Extracted RNA was treated with RNase-free DNase (Qiagen) and stored at −80°C. RNA concentration was determined using a Qubit fluorometer (Invitrogen) and quality was assessed using a Nanodrop One Spectrophotometer (Thermo Fisher Scientific Inc). rRNA depletion and sequencing were performed at the Joint Genome Institute (JGI; Berkeley, California, USA). From 100 ng of RNA, rRNA was depleted using three QIAseq FastSelect kits (Qiagen): 5S/16S/23S, rRNA Plant, and rRNA Yeast. One heavily degraded sample was discarded. Then, paired-end 2 x 151 bp libraries were sequenced for the remaining 47 samples on an Illumina NovaSeq platform. Generated RNA sequences were prepared for metatranscriptomic analyses using the JGI Integrated Microbial Genomes (IMG) pipeline v.5.1.5 (69) and can be found under the GOLD project ID Gp0612223. IMG assemblies are not included in this analysis, and a more detailed description of sequencing conditions, initial sequence QC, and assembly details can be found in Chuckran et al. 2024 (70)

### Metatranscriptome analysis

Raw reads were downloaded from the JGI genome portal and were QC filtered using bbduk (59) and Sickle (60). BBmap was then used to map QC filtered reads to a concatenated reference MAG file (minid=0.95). Counts of reads mapping to DRAM-annotated genes were identified using featureCounts (71). Expression and normalization counts of mapped transcripts for each annotated gene (excluding rRNA genes) were generated using DESeq2 (72) using default parameters. Differential expression was calculated as the log_2_-fold change compared to the gene expression of the dry soil control group.

### Genomic traits

Genomic traits were calculated using custom scripts written in Python (v 3.8.2)—using the packages *pandas* (73) and NumPy (74); they can be found at https://github.com/PChuckran/Wet_up_traits. To capture genome size for each MAG, we corrected the measured genome size with the completeness of the bin—for medium to high quality MAGs (> 70% complete) we divided the total assembled base pairs by genome completeness as a fraction. For each gene in each MAG we calculated the effective number of codons (ENC’) as described by Novembre 2002 (75)—which uses background nucleotide frequencies to assess levels of codon bias. Lower ENC’ values in a gene represent a fewer number of unique codons used in that gene, and therefore higher codon bias.

To determine the relative bias of ribosomal protein genes, we calculated ΔENC’ (26):

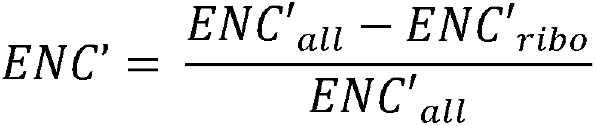

where ENC’_all_ is the effective number of codons for the whole genome, and ENC’_ribo_ is the effective number of codons of the ribosomal protein genes. These ΔENC’ values represent the degree of bias in ribosomal protein genes relative to the rest of the genome, where higher ΔENC’ values represent greater relative codon bias.

Nucleotide frequencies and skews were determined at synonymous and nonsynonymous substitution sites of ribosomal protein genes according to nucleotide degeneracy detailed in Chen et al. 2016 (42). Synonymous site nucleotide frequencies were calculated from nucleotide frequencies at fourfold degenerate sites (sites where any substitution at that location results in the same amino acid), and nonsynonymous frequencies from sites where any substitution would change the encoded amino acid. GC and AT skew were calculated as:

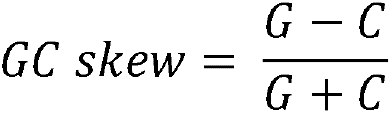

And:

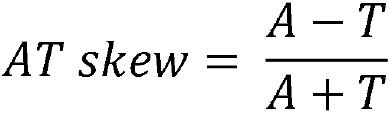

The cost of amino acid synthesis—measured as the total number of phosphate bonds used for synthesis—was derived from Akashi & Gojobori 2002 (76).

### Analysis and model selection

All statistical analyses were conducted in R version 4.2.1 (77) using the tidyverse package (78) and visualized with ggplot2 (79). MAGs were grouped by transcriptional response according to when each MAG was most transcriptionally active, as measured by the total percentage of expressed genes which were upregulated:

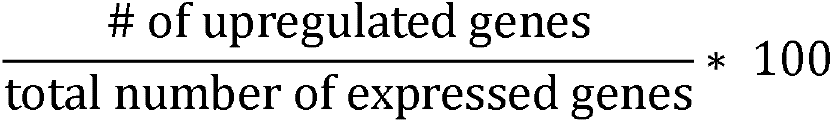

We divided these responses into 4 groups: early responders (most active 3 h post wet-up), middle responders (most active 24, 48, 72 h), late responders (168 h), and sensitive (down-regulated post wet-up; Fig. S1). Differences in genomic traits between transcriptional response groups were determined using an analysis of variance (ANOVA). Tukey’s HSD was used for pairwise comparison between groups. Relationships between genomic traits and AFE values were determined using multiple linear regression. Since we had several related measures of codon bias, to reduce the number of redundant variables, we ran an initial set of model comparisons to assess which codon bias metric best predicted growth rate. Model comparison was conducted using Akaike information criterion (AIC) (80)—with a threshold of -Δ 4, indicating an improvement of a model with the addition of a parameter.

## RESULTS AND DISCUSSION

The rewetting of dry soil represents a reoccurring and impactful change to the soil microbial environment. The response of microbial taxa to this change may be in part determined by traits; however, the influence of the genomic traits commonly associated with growth and transcription on the response to wet-up has yet to be determined. In this study, we re-wet dry soils in a laboratory incubation to create a time-series of the genomic and transcriptomic composition of soils in response to rewetting. To link traits associated with growth and activity during rewetting, we assessed the relationship between genomic traits of metagenome assembled genomes (MAGs) and the affiliated populations’ *in situ* growth rates (measured via MAG targeted quantitative stable isotope probing; qSIP), and transcriptional responses (as measured through metatranscriptomics).

### Traits associated with growth

We assessed the relationship between growth rate and several genome-level and ribosomal protein gene-level traits. This included: codon usage bias, GC content, amino acid synthesis cost (∼P bonds used in synthesis), average amino acid C:N, AT and GC-skew, and genome size. We used atom fraction excess (AFE) values as the response variable, with isotopic enrichment being used as index of growth. The best model (indicated through AIC and R^2^ values) predicted growth using the codon bias in ribosomal protein genes (measured as the effective number of codons; ENC_ribo_), ribosomal protein GC content, and genome size (R^2^ = 0.45, Table S1a&b).

Amongst these traits, ENC’_ribo_ predicted growth better than estimated genome size, or ribosomal GC content (Fig. 1, Table S1A). This demonstrates that codon bias can be predictive of growth not only in pure cultures, as previously established (26), but also in complex soil microbial communities responding to a natural pulse phenomenon. Although there has been no analysis observing the broad-scale distribution of codon bias in soils, previous work suggests bacteria in soils with lower C, higher pH, and less rainfall tended to preferentially use one synonymous codon over another (81). High codon bias may be an important attribute for soil bacteria experiencing transient water and nutrient availability, enabling quicker responses to sudden pulse events. This highlights the importance of codon bias as a critical trait when considering the adaptation of soil bacteria to changing precipitation patterns in the context of climate change.

**Figure 1.**
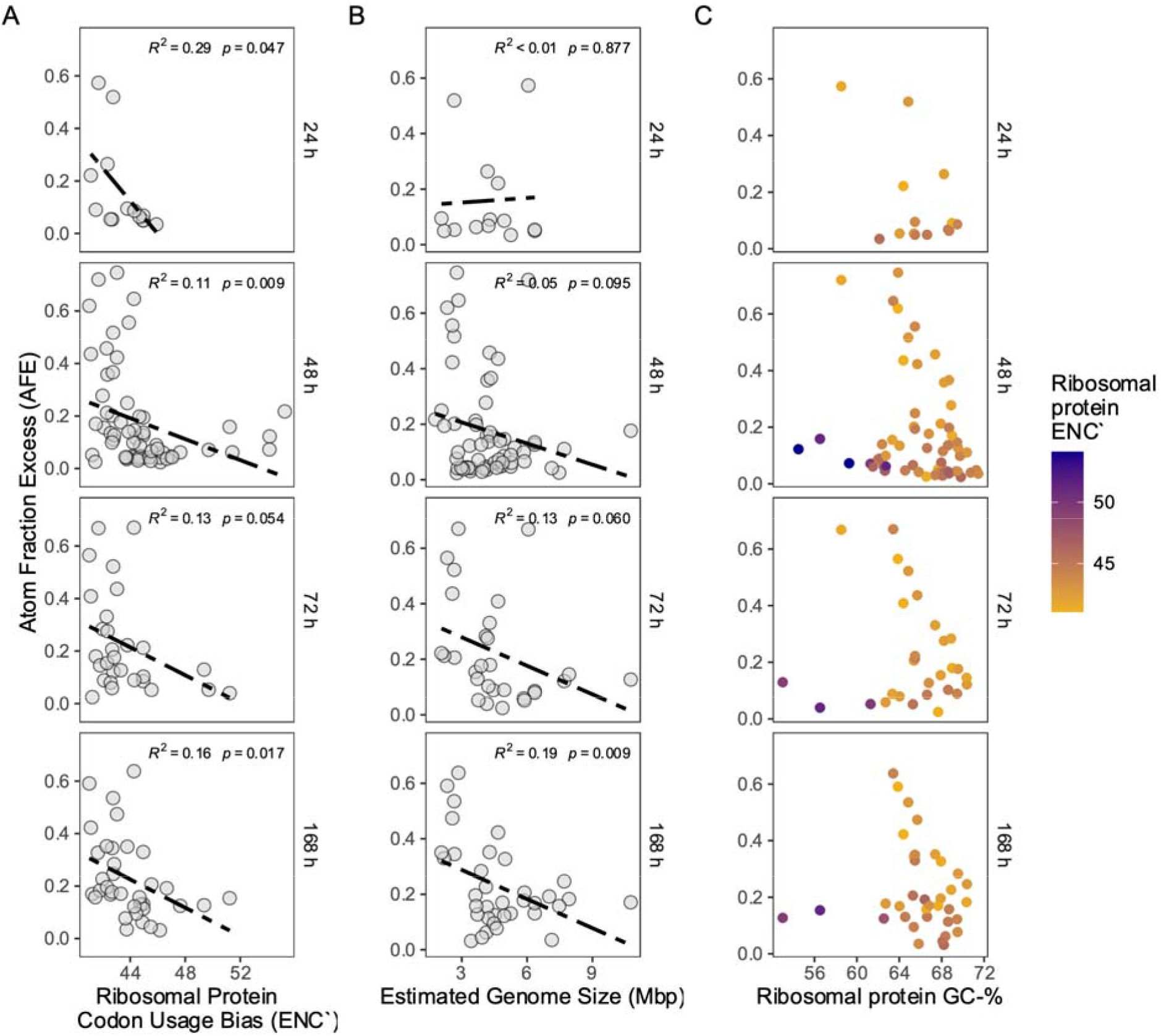
Effect of a 1 week H_2_^18^O wet-up experiment of annual grassland soils from Hopland, CA on bacterial growth and genomic traits. **(A)** Relationships between taxon-specific growth (atom fraction excess (AFE) of ^18^O) and effective number of codons in ribosomal protein genes (ENC) and **(B**) genome size (Mbp, derived from metagenome assembled genomes (MAGs)), for samples collected at multiple post wet-up timepointsan, and **(C)** versus GC content of ribosomal protein genes post wet-up (the time where the most MAGs were represented), with color indicating ENC values. Lower ENC values indicate a higher level of codon bias.

Ribosomal GC content was negatively correlated with growth rate (Table S1, Fig. 1C)— provided that codon bias was high (i.e. ENC_ribo_ was low, Fig. 1C, Fig. S2). Although what drives this relationship is unclear, these results suggest that a lower GC content poses an advantage for rapid growth in response to wet-up. We hypothesize that given the greater metabolic (ATP) cost of the GC base pair compared to the AT base pair (42), higher GC content could slow growth rate. GC content is also associated with metabolic strategy, where lower GC content is typically associated with greater preference for glycolytic carbon sources, such as sugars and compounds with a higher carbon to nitrogen ratio (C:N) (82). It’s possible that the sudden increase in bioavailable carbohydrates upon rewetting (83) gives an advantage to lower GC bacteria. This trend is particularly notable considering that genomic GC-content is generally predictive of codon bias (30), where more extreme GC values will naturally result in higher levels of codon bias (an example of this relationship with the MAGs used in this study is shown in Fig. S3). In this way, the emergence of neutral GC content and high codon bias requires that each of these traits be selected for independently in taxa with these traits.

We also found that smaller genomes (< 3 Mbp) were associated with higher growth rates at later time points, particularly at 168 h post wet-up (Fig. 1B)—where a 75% reduction in genome size resulted in ∼200% increase in growth rate. Smaller genomes may reduce the cost of replication, a key factor in responding to sudden changes in environmental conditions. As a corollary, large genomes that enable metabolic flexibility may come at the expense of growth rate (45). This may also explain why smaller genomes are more prevalent in arid and carbon poor soils (81, 84, 85), since the ability to quickly respond to the momentary availability of water and resources may be fundamental to survival and persistence.

Our results contrast with observations from marine systems, where free-living bacteria with small streamlined genomes (1-2 Mbp) tend to have slower growth rates (43). This biome-scale inconsistency between growth rate and genome size may reflect fundamentally different life-strategies that both leverage simplified genomes. In marine environments, reduced genomic complexity and a high surface area to volume ratio occur in response to nutrient limitation— reducing the total cost of replication while increasing the likelihood of capturing dissolved nutrients (43). However, these traits in marine environments are typically not associated with faster growth rates (18). In contrast, for soil communities responding to a rapid pulse of resources or change in environmental conditions, the lower costs associated with replicating smaller genomes may allow for faster growth in response to flux in nutrient availability. The reduced cost of smaller genomes might serve both strategies well, and the relationship between genome size and growth rate could be environment specific.

### Traits associated with transcription

To assess the influence of genomic traits on transcription, we separated our dataset into 4 temporal categories according to peak transcriptional responses—based on the proportion of expressed genes that were significantly upregulated at each timepoint (see methods, Fig. S1).

High codon bias, in addition to being a predictor of growth, has also been shown to increase gene expression (36, 37). We found that the codon bias of ribosomal protein genes was significantly different between transcriptional response group (ANOVA; *p* < 0.01), with a pairwise comparison (Tukeys HSD, *p* < 0.05) indicating the exception being between sensitive (down- regulated post wet-up) and middle or late-responding taxa. Codon bias corresponded with transcription timing, where early responding taxa had a significantly higher level of codon bias than all other groups, followed by middle-responding taxa, and then late responding and sensitive taxa (Fig 2A). Regulation of ribosomal protein genes corresponded to transcriptional response group (Fig. S4), and although this is not a particularly surprising result considering response categories were curated, it does demonstrate a relationship between codon bias and transcriptional regulation of individual sets of genes. Although the importance of codon bias for transcription has been shown for individual taxa (36, 37), this result elucidates that codon usage may be an important feature in dictating the response of bacteria living in complex soil communities. Further, this result demonstrates that codon usage is not only important for the growth response of bacteria during rewetting, but also plays a role in the timing of transcriptional responses.

**Figure 2:**
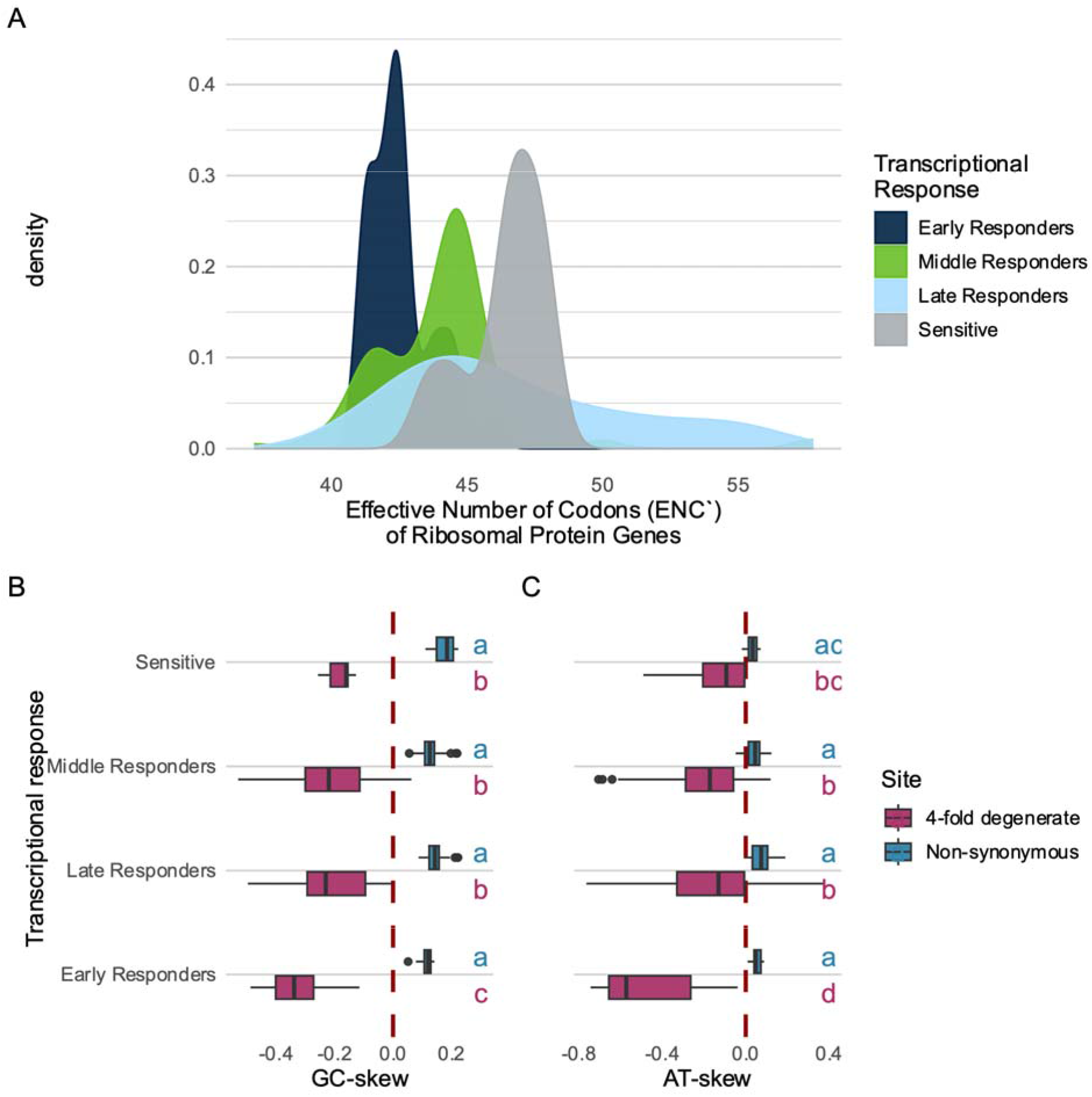
Measures of ribosomal protein response to soil wet-up in a grassland soil. Codon usage bias of ribosomal protein genes colored by transcriptional response type **(A)**. The synonymous and non-synonymous GC (**B**) and AT **(C)** skew by transcriptional response, with color indicating nucleotide site-type. Four-fold degenerate sites describe a type of synonymous substitution site where all possible nucleotide substitutions would result in the same amino acid.

Nucleotide selection, in addition to its inherent relationship with codon usage, can influence transcription due to the cost of nucleotide synthesis. Each nucleotide requires a different amount of energy to synthesize, where A > T, G > C, and G+C > A+T (42). At synonymous sites, where nucleotide substitutions do not change the encoded amino acid, nucleotides which are cheaper to synthesize have been shown to be associated with faster rates of transcription (42). We found that early responding organisms displayed a lower AT (A-T/A+T) and GC skew (G-C/G+C) at synonymous sites for ribosomal protein genes (Fig. 2 b&c). Our findings align with results from Chen et al. (2016) (42), which demonstrated that the lower cost of U and C at synonymous sites was associated with increased gene transcription. No such effect was observed at nonsynonymous sites (Fig. 2 b&c), showing that cheaper nucleotide selection is maintained only when peptide sequence is not impacted. This distinction between synonymous and non-synonymous sites extends to all transcriptional response categories evidencing the cost-saving encoding of lower GC-and AT-skew only when amino acid sequence is not compromised (Fig. 2 b&c). Notably, GC skew and AT skew of synonymous substitutions on ribosomal protein genes were weakly negatively correlated to growth rate at certain time points (Fig. S5A). We did find a relationship between nonsynonymous nucleotide skew and growth (Fig. S5 C&D), which we discuss in the Supplemental Results and Discussion. These correlations are weakly significant in the context of other traits, but suggest that these cost-saving traits may be associated with fast-growing taxa. These results underscore the disparity between traits affecting transcription and those having a more pronounced influence on growth.

### The relationship between growth and transcription

Transcription, although necessary for growth, has been argued to be a poor indicator of activity (28). Translational efficiency may additionally influence growth rate, and while we cannot directly measure the contribution of higher transcription vs translational efficiency on growth in these data, we can assess the relationship between transcriptional response and growth. We did not find a strong relationship between the overall expression of ribosomal protein genes and growth rate, as measured by AFE (Fig. 3a), nor did we find a significant relationship between transcriptional response group and growth rate (Fig. 3b). These results demonstrate that although transcription may be necessary for growth, levels of RNA may not serve as a reliable quantitative proxy for growth rate at the community scale, and corroborates prior work indicating rRNA serves as a poor metric for activity (28).

**Figure 3:**
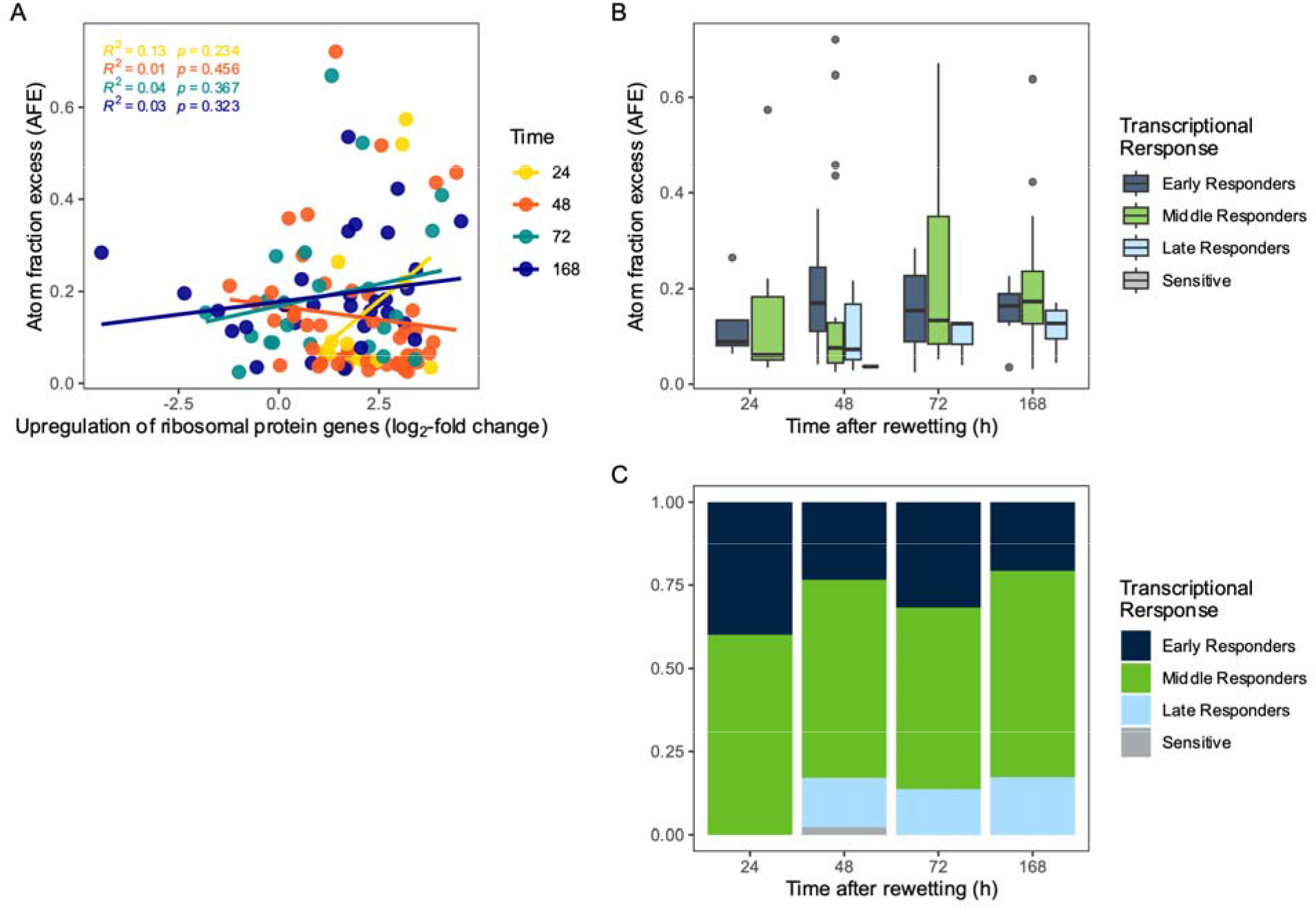
The relationship between growth (as indicated through Atom Fraction Excess, AFE) and transcription after rewetting for metagenome assembled genomes (MAGs). The level of upregulation of ribosomal protein genes in each MAG versus growth (AFE), with color indicating time **(A)**. AFE of each transcriptional response group over time (**B**), and the proportion of growing MAGs that could be assigned to a transcriptional response group at each time point (**C**).

We next explored whether transcriptional response patterns corresponded with growth rate. We found that the transcriptionally most active early were more likely to be growing early, and taxa that were most transcriptionally active later were more likely to be growing later (Fig. 3C). This connection between transcriptional activity and growth does suggest that even if transcription may not be a good predictor or proxy for growth, the timing of transcription generally follows the timing of growth after rewetting.

### Phylogenetic conservation of trait relationships

Genomic traits such as genome size (86), codon usage, and nucleotide selection are phylogenetically conserved—raising the question as to the relative impact of these traits within the context of taxonomy. For example, early growing taxa were dominated by Proteobacteria (37-57% of growing taxa) (Fig. 4A), which often had higher levels of ribosomal codon bias (Fig. 4B, Fig. S7A). This would suggest that the predictive power of genomic traits are phylogenetically conserved. The phylogenetic distribution of these traits may explain results from other rewetting studies, which have found a high abundance of Proteobacteria in response to wet-up (58, 87).

**Figure 4:**
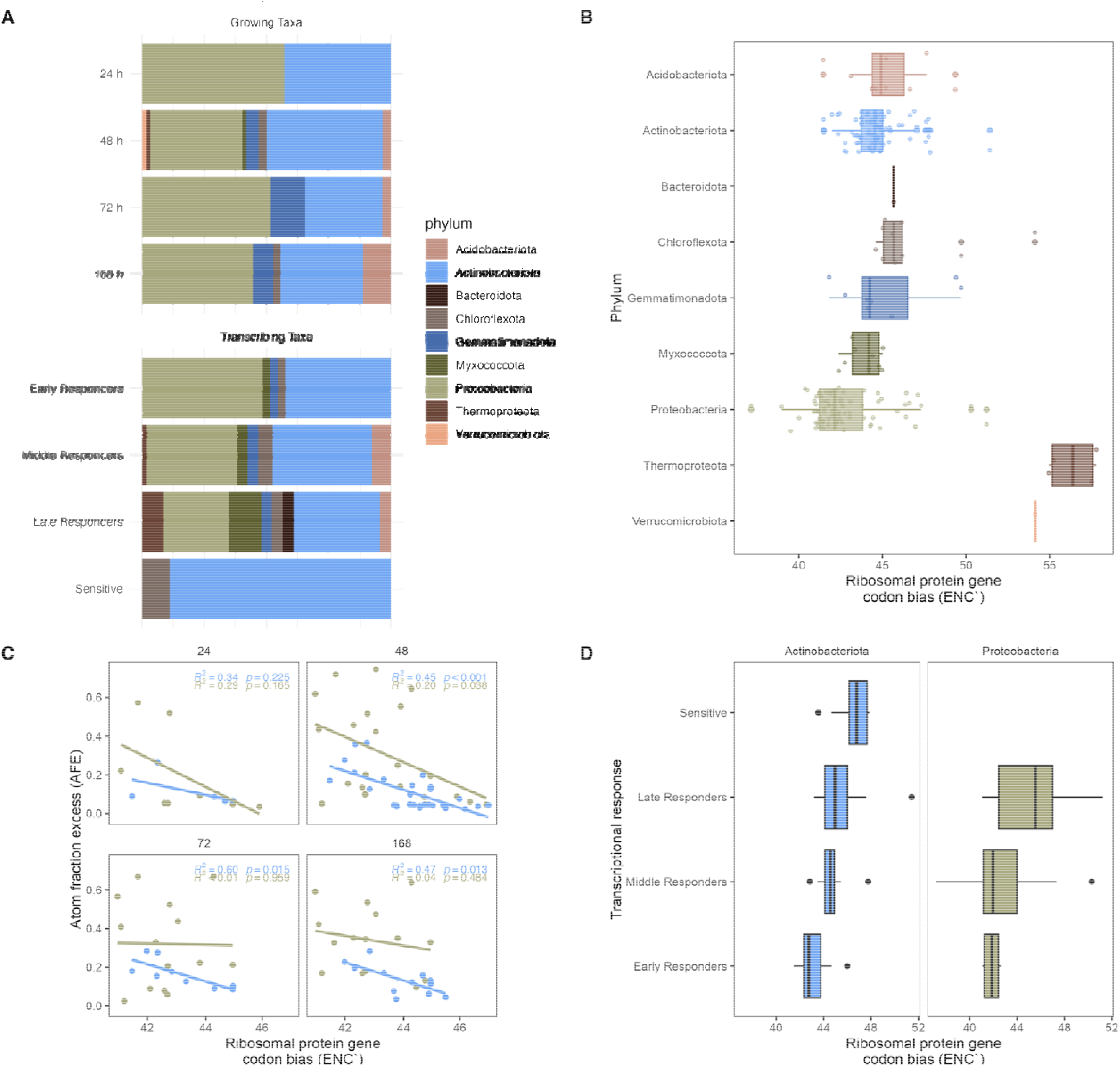
Taxonomic breakdown of metagenome assembled genomes (MAGs) by phylum growing at each time point and by transcriptional response category (**A**). Ribosomal protein gene codon usage bias (ENC_ribo_) for all growing or transcribing MAGs separated by phylum (**B**). Atom fraction excess, and index of growth, versus ribosomal protein codon usage bias (ENC), separated by the two most represented phylum, Actinobacteria and Proteobacteria (**C**). Ribosomal protein codon usage bias (ENC) by transcriptional response category and phylum (**D**).

However, the observed relationships between traits and activity could also exist within a taxonomic group—meaning that the influence of genomic traits on the rewetting response would be more generalized features of soil bacteria. The number of MAGs recovered in this study does not allow for a comprehensive quantitative analysis of the contribution of phylogeny vs traits to activity; however, we can assess whether patterns between traits and growth and transcription emerge withing a phylogenetic group. We chose to analyze the relationships between growth, transcription, and traits among the two most abundant and prevalent phyla at HREC, Proteobacteria and Actinobacteria (Fig. 4A). Consistent with total community observations, higher AFE values tended to correlate with higher levels of codon bias (Fig. 4C). We also found that transcriptional response categories followed a similar trend to community level observations, with early transcriptional response being associated with higher levels of codon bias (Fig. 4D).

Although we cannot properly assess the influence of other genomic traits in a complex model due to sample size constraints, we did find some notable trends when separated by phylum: genome size in Proteobacteria was significantly negatively related to growth at 48 and 168 h (Fig. S7B), and nucleotide skew followed community-level trends with respect to transcriptional response category (Fig. S7 C&D). These results suggest that relationships between growth or transcription with genomic traits are not strictly reliant on phylum-level taxonomy and may represent fundamental trade-offs for soil bacteria.

In contrast, we did not find a relationship between ribosomal GC-content and growth on the phylum level. Rather, we found this relationship in two fast-growing families of Proteobacteria: *Sphingomonadacea* and *Burkholderiacea* (Fig. S7E). This indicates that low GC-content is not a trait universally related to growth rate in soil bacterial taxa, but that this is a phylogenetically conserved relationship. Previous studies observed a high representation of *Sphingomonadacea* in growing bacteria post rewetting (87, 88) and these genomic trait relationships might be related to a unique life strategy posing a fitness advantage—perhaps due to substrate preference, which has been shown to be related to GC content (82).

## Conclusion

Rapid environmental change such as rewetting may play an important role in the selection of traits in soil bacteria. Using a combination of metagenome-assembled genomes, transcriptomics, and ^18^O-water qSIP, we evaluated genomic traits associated with transcriptional activity and growth of soil microorganisms in the period immediately following soil rewetting. Growth was not proportionally related to levels transcription; however, timing of transcription and growth were generally coupled. We found that transcription was associated with nucleotide synthesis cost as well as codon usage bias, and that growth was associated with high codon usage bias, lower GC content, and smaller genome size. This work highlights the importance of genomic traits to short-term responses in systems characterized by pulses of water and nutrient availability, enhancing our understanding of metrics for assessing the functional potential of soil bacteria and offering important perspective on the broad scale ecological distribution of genomic traits in soil.

## Data Availability

Metatranscriptome data are located on the NCBI database, IMG/M database, and JGI genome portal. Sample names, links to sequence data, and descriptions can be found in the data release (https://doi.org/10.1128/mra.00322-24) (70). MAGs can be accessed from the NCBI database under the BioProject PRJNA856348 and PRJNA718849.

## Acknowledgements

We would like to thank Mengting Maggie Yuan, Erin Nuccio, Aaron Chew, Anne Kakouridis, Kate Zhalnina, Nameer Baker, Rachel Hestrin, Caleb Herman, Javier Ceja-Navarro, Christina Ramon, Alex Greenlon, Ryan Gini, Ilexis Chu-Jacoby, Melissa Lafler, Keith Morrison, Amrita Bhattacharyya, Heejung Cho, Angela Hodge, Albert Molina, Don Herman, and Xiao Bin Max Li for their help with the experimental set-up and sample processing. We thank Alex Greenlon, Jillian Banfield, and Rohan Sachdeva for their assistance with MAG creation and bioinformatic expertise, and Carl Roybal for intellectual contributions and manuscript proofing.

This research was supported by the U.S. Department of Energy (DOE), Office of Biological and Environmental Research (BER), Genomic Science Program Lawrence Livermore National Laboratory (LLNL) ‘Microbes Persist’ Soil Microbiome Scientific Focus Area SCW1632, and a subaward to UC Berkeley. Field plots and precipitation management were initially generated via DOE BER awards DE-SC0020163 and DE-SC0016247 (to MKF) and awards SCW1589, and SCW1421 (to JPR). Sequencing was conducted at the Joint Genome Institute (JGI) via a Department of Energy Biological and Environmental Support Science award #508594 (to JPR). Research Work at Lawrence Livermore National Laboratory was conducted under the auspices of the U.S. DOE under contract DE-AC52-07NA27344.

