## Supplemental Material for "Codon bias, nucleotide selection, and genome size predict *in situ* bacterial growth rate and transcription in rewetted soil"

Supplemental Results and Discussion:

We found a slight positive relationship between nonsynonymous GC-skew and growth (Fig. S5C), and a surprisingly positive relationship between nonsynonymous AT-skew and growth (Fig. S5D, R^2^ = 0.22, p < 0.01). This could potentially reflect the inverse relationship between nucleotide and amino acid cost. More expensive nucleotides tend to encode less energetically expensive amino acids (40), and this relationship could reflect this cost conservation. However, amino acid cost was not associated with growth (Fig. S6). When we examined the amino acid composition as related to nonsynonymous substitutions, we found that AT skew most closely correlated with the abundance of lysine (encoded for by AAG and AAA)—yet it is unclear what specifically about lysine would be causing this relationship. Lysine is not notably cheaper to synthesize and has elemental ratios similar to other amino acids. This relationship may be due to features of the ribosomal protein structure, the analysis of which is beyond the scope of this study. Alternatively, the relationship could also be caused by correlations between AT skew and other genomic features, especially GC content (linear regression R^2^= 0.46). Further, the addition of nonsynonymous AT skew to the model outlined in Supplemental Table 1 did not explain much more variation compared to the base model—suggesting a potentially spurious correlation.

**Supplemental Table 1 a:**

| Model | df | AIC |
| --- | --- | --- |
| ENC`ribo | 3 | -104.55531 |
| deltaENC | 3 | -119.64409 |
| Non-synonymous AT-skew | 3 | -113.61949 |
| ENC` | 3 | -81.09837 |
| Genome size (bp) | 3 | -91.71022 |
| Ribosomal protein GC content (%) | 3 | -87.20576 |
| Genome GC content (%) | 3 | -84.24045 |
| Non-synonymous AT-skew X deltaENC | 4 | -142.59544 |
| deltaENC X Ribiosomal Protein GC | 4 | -122.94479 |
| deltaENC X Genome size | 4 | -130.15463 |
| ENC`ribo X Genome size (bp) x Ribosomal protein GC | 5 | -170.21837 |
| deltaENC X Genome size (bp) x Ribosomal protein GC | 5 | -132.30391 |
| deltaENC X Ribiosomal Protein GC X Non-synonymous AT-skew | 5 | -133.69058 |
| deltaENC X Genome size X Non-synonymous AT-skew | 5 | -144.64803 |
| ENC`ribo X Genome size (bp) x Ribosomal protein GC X Non-synonymous AT-skew | 6 | -174.90644 |

**Supplemental Table 1 b:**

| Coefficients: |  |  |  |  |  |
| --- | --- | --- | --- | --- | --- |
|  | Estimate | Std. Error | t value | p-value |  |
| (Intercept) | 4.23E+00 | 3.87E-01 | 10.907 | < 2e-16 | *** |
| ENC`ribo | -4.90E-02 | 4.93E-03 | -9.932 | < 2e-16 | *** |
| Genome size (bp) | -3.21E-08 | 6.43E-09 | -4.984 | 1.74E-06 | *** |
| Ribosomal Protein GC (%) | -2.61E+00 | 3.41E-01 | -7.653 | 2.45E-12 | *** |
| Signif. codes: 0 ‘***’ 0.001 ‘**’ 0.01 ‘*’ 0.05 ‘.’ 0.1 ‘ ’ 1 | | | |  |  |
| Residual standard error: 0.1345 on 146 degrees of freedom | | | |  |  |
| Multiple R-squared: 0.4631, | Adjusted R-squared: 0.4521 | | |  |  |
| F-statistic: 41.98 on 3 and 146 DF, p-value: < 2.2e-16 | | |  |  |  |

**Supplemental Table 1 c:**

| Coefficients: |  |  |  |  |  |
| --- | --- | --- | --- | --- | --- |
|  | Estimate | Std. Error | t value | p-value |  |
| (Intercept) | 3.52E+00 | 5.17E-01 | 6.82 | 2.42E-10 | *** |
| Non-synonymous AT skew | 1.00E+00 | 4.44E-01 | 2.25 | 0.026 | * |
| ENC`ribo | -4.48E-02 | 5.51E-03 | -8.131 | 1.91E-13 | *** |
| Genome Size (bp) | -3.05E-08 | 6.27E-09 | -4.866 | 2.99E-06 | *** |
| Ribosomal Protein GC content (%) | -1.92E+00 | 4.54E-01 | -4.217 | 4.39E-05 | *** |
| Signif. codes: 0 ‘***’ 0.001 ‘**’ 0.01 ‘*’ 0.05 ‘.’ 0.1 ‘ ’ 1 | | |  |  |  |
| Residual standard error: 0.1304 on 142 degrees of freedom | | |  |  |  |
| Multiple R-squared: 0.5036, | Adjusted R-squared: 0.4896 | | |  |  |
| F-statistic: 36.01 on 4 and 142 DF, p-value: < 2.2e-16 | | |  |  |  |

**Supplemental Figure 1:**

Examples of gene regulation patterns for each transcriptional response category. MAGs were grouped into the following categories based on the net proportion of significantly upregulated genes over time after rewetting.

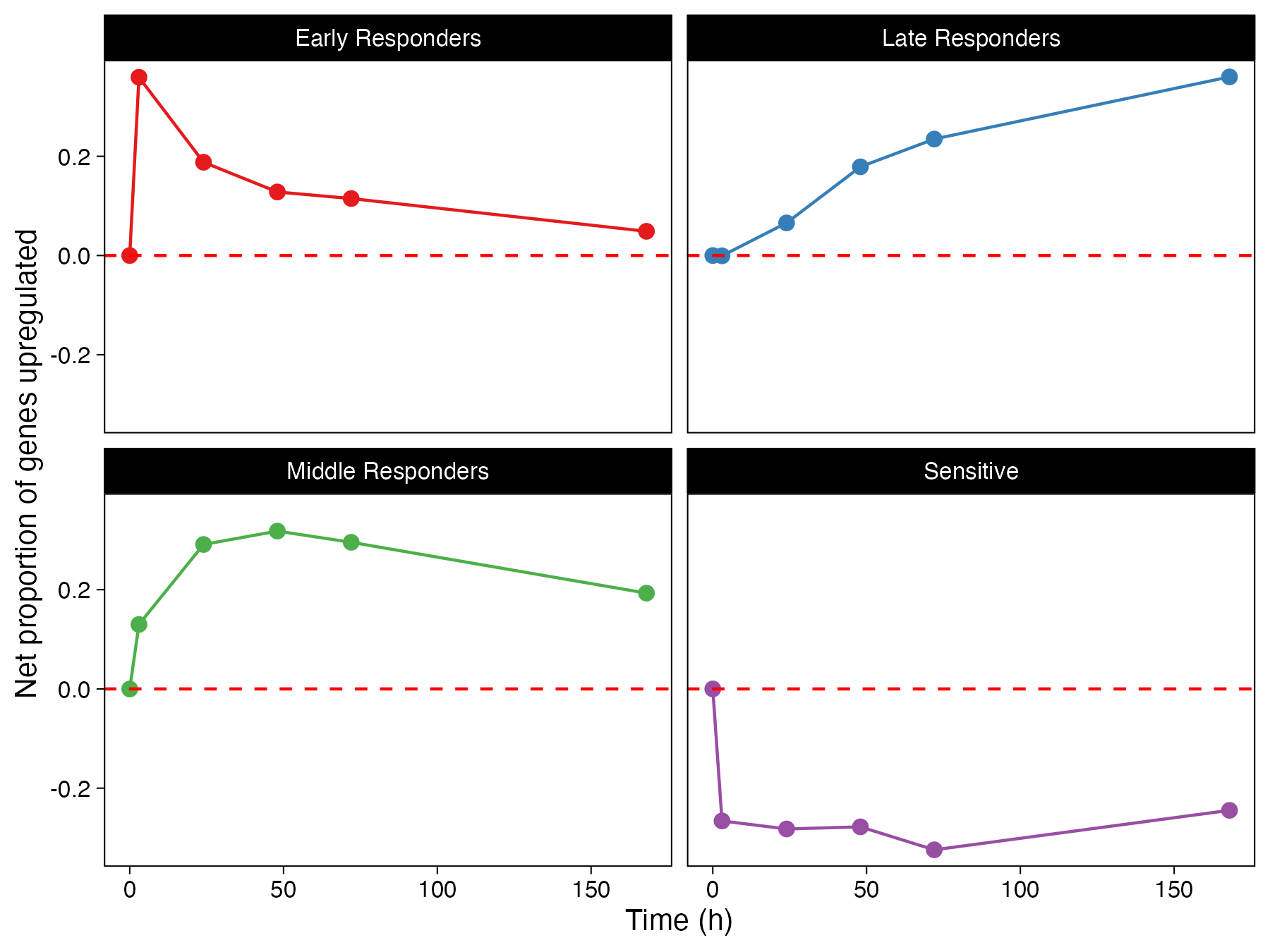

**Supplemental Figure 2**

Predicted relationship between ribosomal protein gene GC content and growth (as measured through atom fraction excess; AFE), with color indicating how this relationship varies with ribosomal protein gene codon bias based on the predictions from our model.

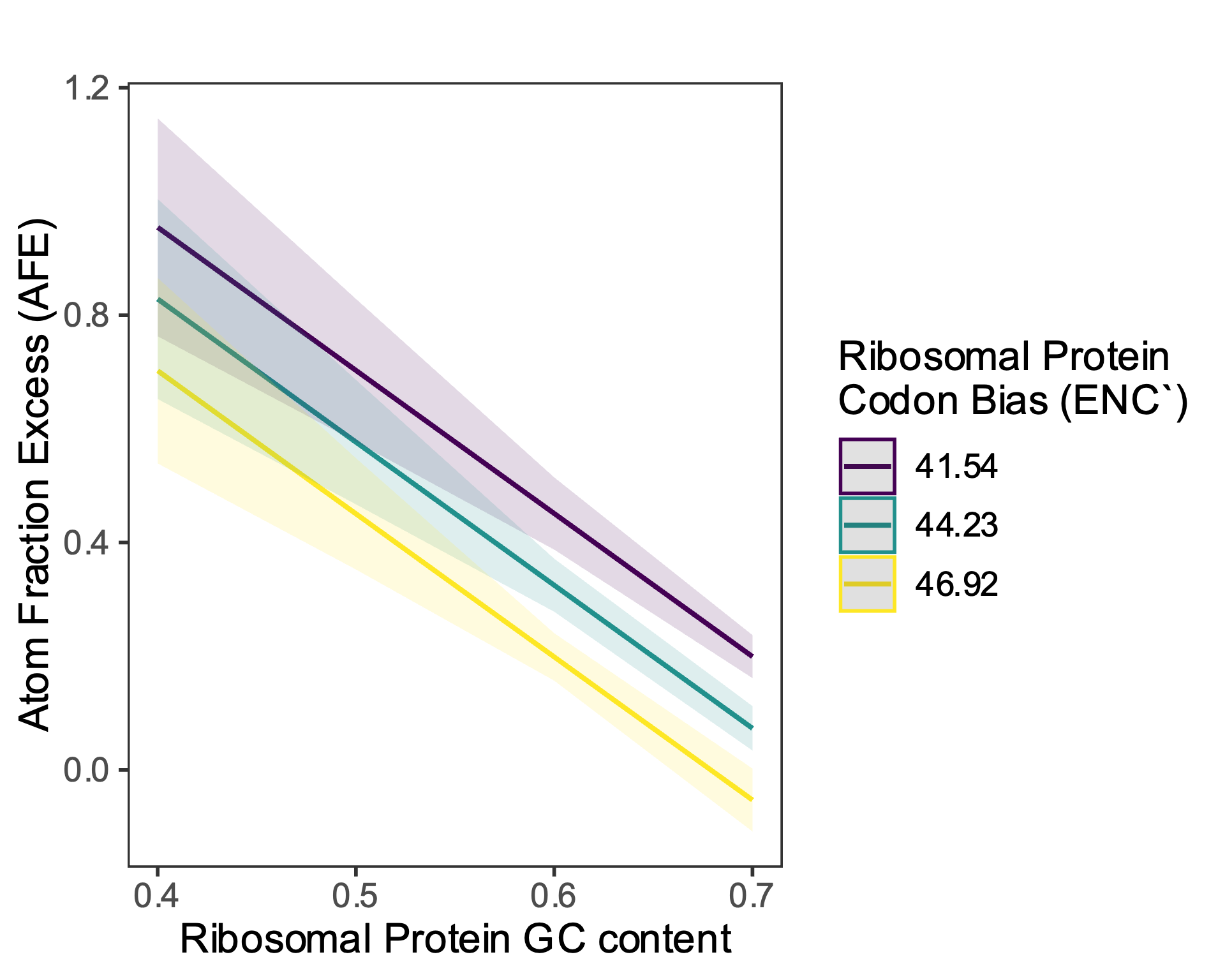

**Supplemental Figure 3**

Codon usage bias (ENC`) as it relates to GC content, for the whole genome (left) and for ribosomal protein genes (right)

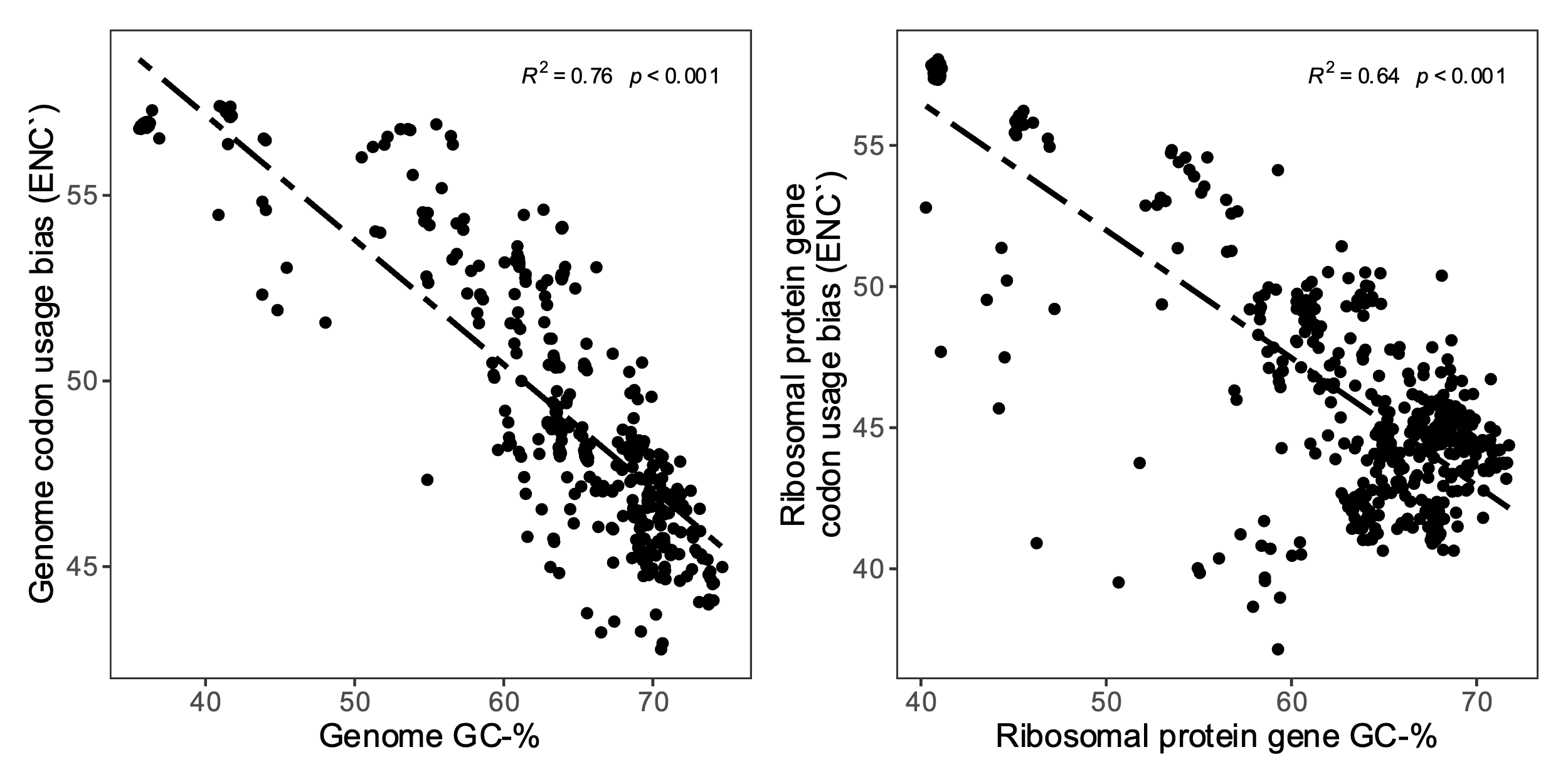

**Supplemental Figure 4**

Mean upregulation of ribosomal protein genes after rewetting (expressed as log_2_-fold change vs initial timepoint) for each transcriptional response category.

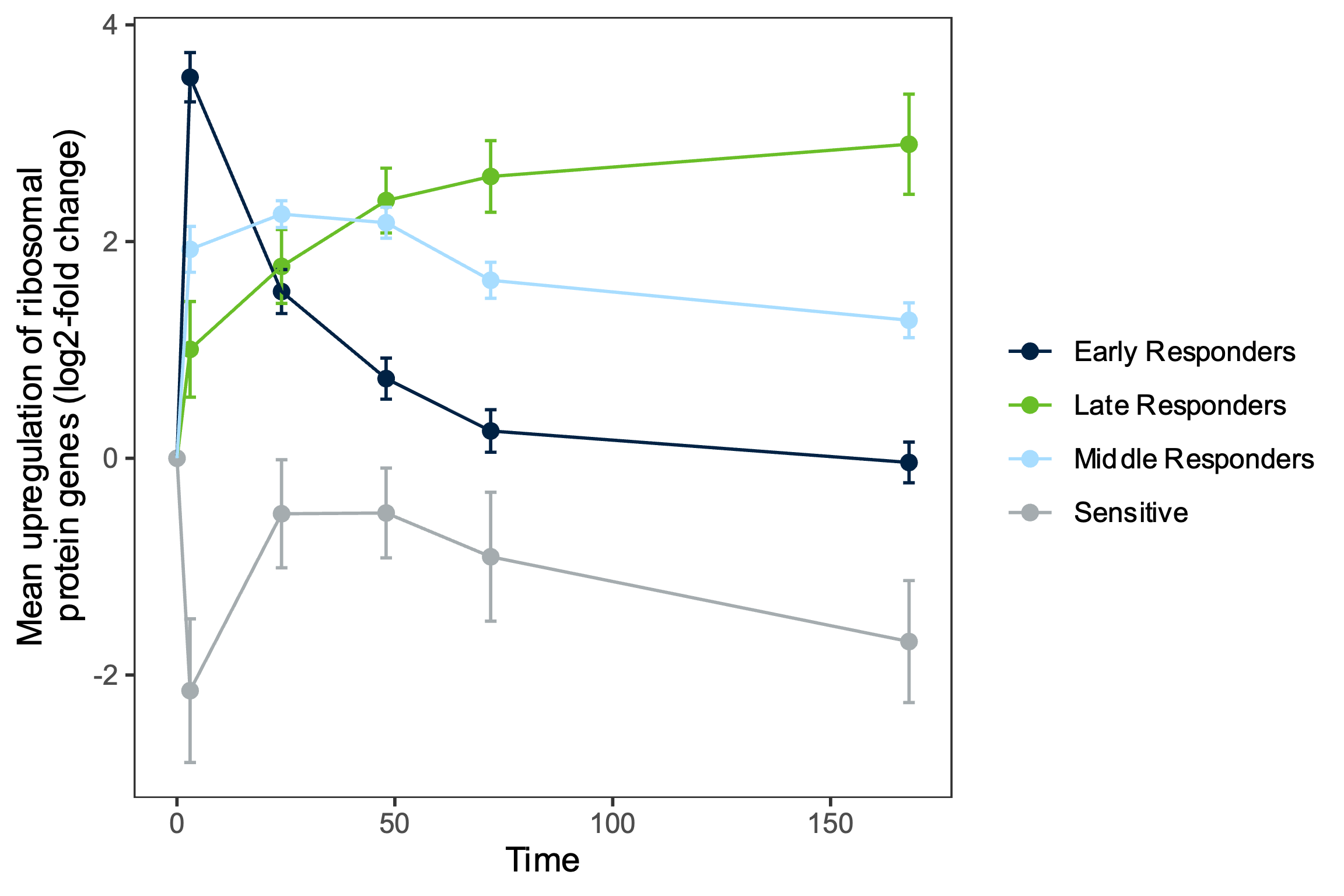

**Supplemental Figure 5**

Nucleotide selection in ribosomal protein genes as it relates to growth rate—as measured through Atom Fraction Excess (AFE) values. Synonymous site GC skew (**A**) and AT skew (**B**), and Non-synonymous site GC skew (**C**) and AT skew (**D**). Relationships are separated by time after rewetting, which is indicated by color.

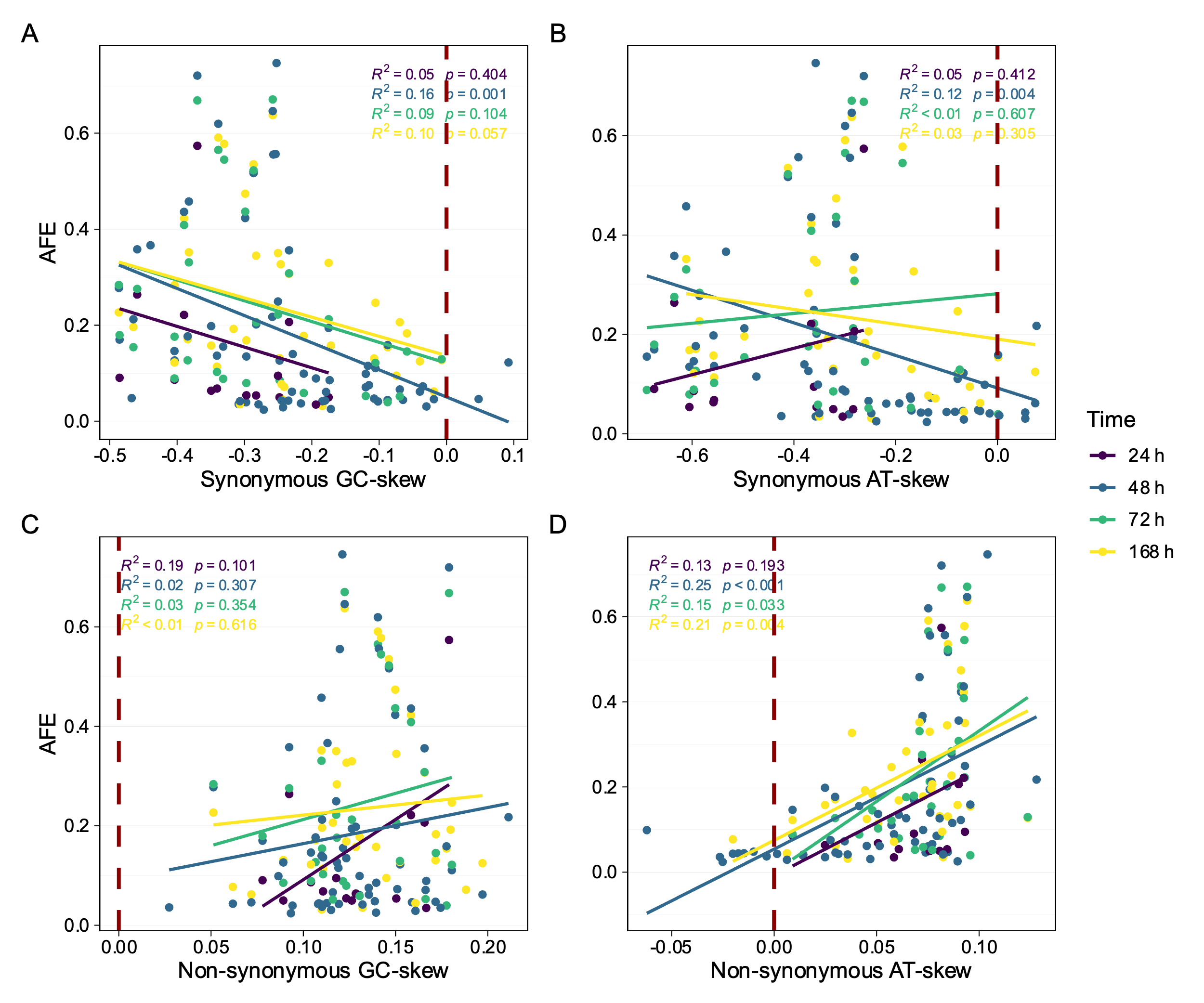

**Supplemental Figure 6**

Growth, as measured through Atom Fraction Excess (AFE) values, as a function of mean synthesis cost of ribosomal protein synthesis, as measured by the mean number of P bonds used in synthesis per amino acid.

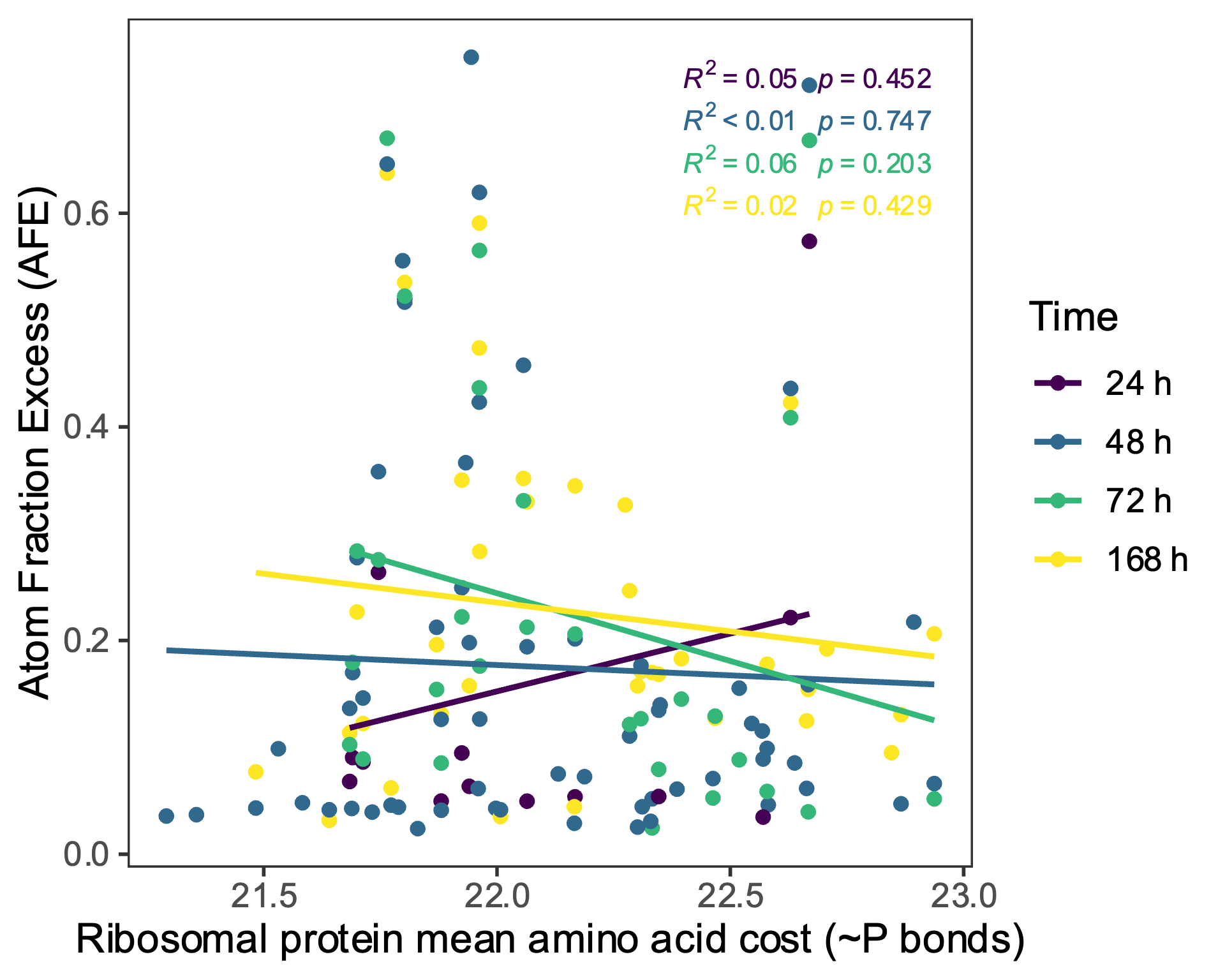

**Supplemental Figure 7**

Select genomic trait relationships in the context of phylogeny. Ribosomal protein gene codon usage bias (ENC`) for all binned MAGs separated by phylum (**A**). For Actinobacteria and Proteobacteria: the relationship between genome size and growth (as measured through atom fraction excess, AFE) (**B**); transcriptional response category and synonymous GC-skew (**C**) and AT-skew (**D**) in ribosomal protein genes. Growth as a function of the GC content of ribosomal protein genes, with *Sphingomonadacea* and *Burkholderiacea* highlighted (**E**).

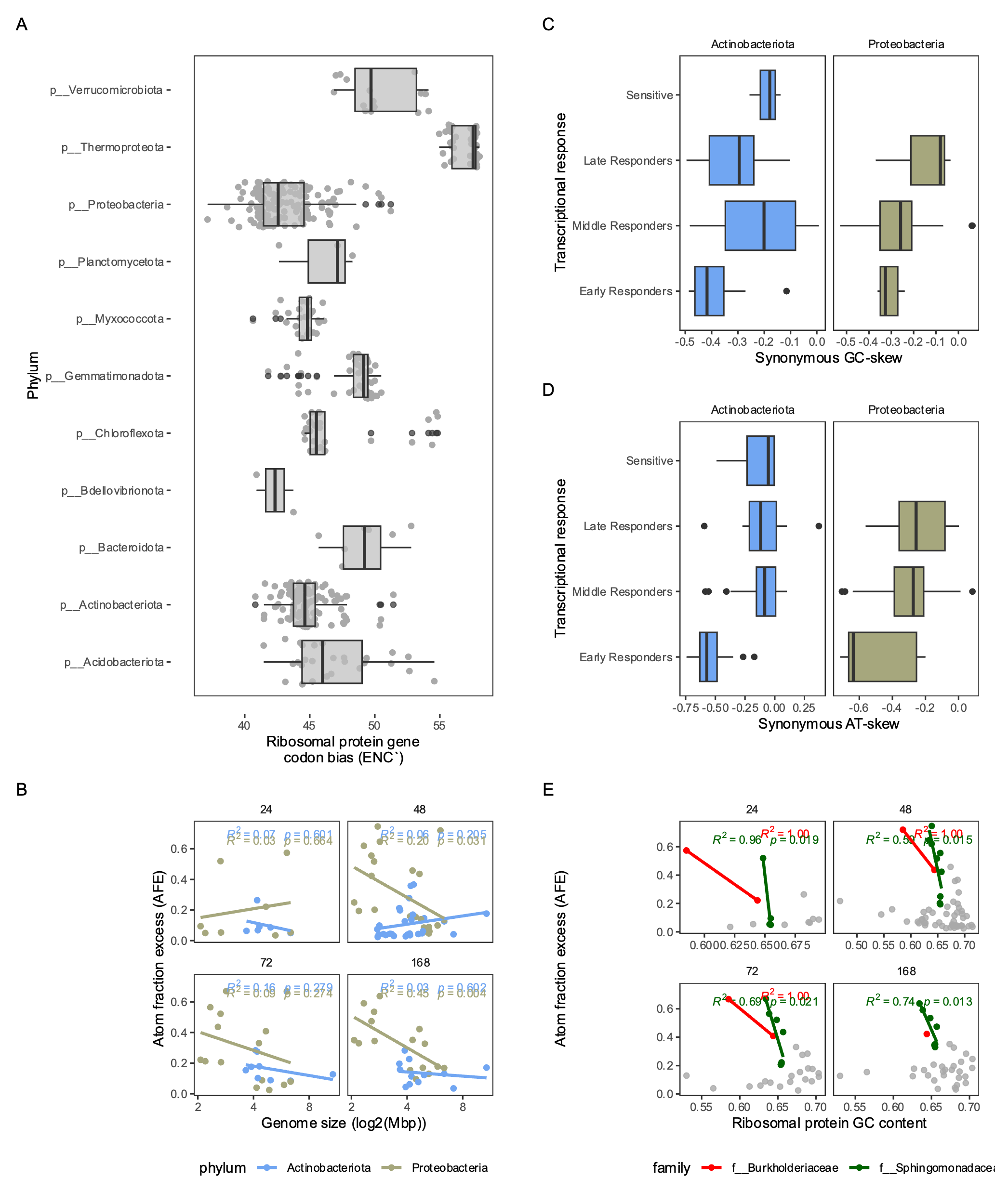
